# Histone deacetylases regulate organ-specific growth in a horned beetle

**DOI:** 10.1101/2023.11.30.569342

**Authors:** Yonggang Hu, Jordan R. Crabtree, Anna L. M. Macagno, Armin P. Moczek

## Abstract

**Background:** Nutrient availability is among the most widespread means by which environmental variability affects developmental outcomes. Because almost all cells within an individual organism share the same genome, structure-specific growth responses must result from changes in gene regulation. Earlier work suggested that *histone deacetylases* (*HDACs*) may serve as epigenetic regulators linking nutritional conditions to trait-specific development. Here we expand on this work by assessing the function of diverse *HDACs* in the structure-specific growth of both sex-shared and sex-specific traits including evolutionarily novel structures in the horned dung beetle *Onthophagus taurus*.

**Results:** We identified five *HDAC* members whose down-regulation yielded highly variable mortality depending on which *HDAC* member was targeted. We then show that *HDAC1*, *3*, and *4* operate in both a gene- and trait-specific manner in the regulation of nutrition-responsiveness of appendage size and shape. Specifically, *HDAC 1, 3,* or *4* knockdown diminished wing size similarly while leg development was differentially affected by RNAi targeting *HDAC3* and *HDAC4*. In addition, depletion of *HDAC3* transcript resulted in a more rounded shape of genitalia at the pupal stage and decreased the length of adult aedeagus across all body sizes. Most importantly, we find that *HDAC3* and *4* pattern the morphology and regulate the scaling of evolutionarily novel head and thoracic horns as a function of nutritional variation.

**Conclusion:** Collectively, our results suggest that both functional overlap and division of labor among *HDAC* members contribute to morphological diversification of both conventional and recently evolved appendages. More generally, our work raises the possibility that *HDAC*-mediated scaling relationships and their evolution may underpin morphological diversification within and across insect species broadly.

## BACKGROUND

Phenotypic plasticity is the ability of an organism to change its phenotype in response to environmental stimuli (Schlichting and Pigliucci 1998; West-Eberhard 2003), a universal phenomenon in the living world. Diverse abiotic (e.g. temperature, photoperiod) and biotic (e.g. conspecific density) may influence growth and differentiation (Régnière et al., 2012; Prudic et al., 2011; O’Leary et al., 2017). Among those, nutrition is one of the most widespread means by which environmental variability affects developmental outcomes. Lack of essential nutrients can slow or arrest development, and sometimes trigger alternative developmental programs, such as the diapause of insects and worms (Lafuente and Beldade 2019). Nutrition also serves as a major determinant of animal size and shape (Andersen et al. 2013; Rohner et al. 2022), with poor nutrition generally yielding reduced growth and final adult body size in animals with determinate growth such as insects and mammals. However, different body parts within an individual typically differ in their response to nutrient availability. For instance, brain size in mammals and male genital size in arthropods are relatively nutrition-insensitive (Koh et al. 2005; Eberhard 2009), whereas secondary sexual traits such as male horns of dung and rhinoceros beetles (Moczek 1998; Ito et al. 2013), or mandibles of stag and broad-horned flour beetles (Gotoh et al. 2014; Okada et al. 2019) are exquisitely sensitive to nutritional variation during development. Such trait-specific scaling relationships therefore contribute in important ways to shape morphological diversity within and among taxa. Here we investigate the role of epigenetic modifications via organ-specific growth responses in an insect model.

Because all cells within an individual organism essentially share the same genome, organ- or structure-specific growth must result from changes in gene regulation. Much recent work has described changes in transcription profiles in response to environmental modifications such as nutrient availability, and has begun to identify key regulators of condition-responsive growth (e.g. *insulin*/*IIS* (Emlen et al. 2012; Snell-Rood and Moczek 2012; Xu et al. 2015), *doublesex* (Kijimoto et al. 2012), and *hedgehog* (Kijimoto and Moczek 2016)). However, how such variation in gene expression is achieved in the first place, and then subsequently transduced into organ-specific growth is much less well understood. Here we investigate the role of epigenetic modifications in enabling organ-specific growth, with a particular emphasis on histone modifications. Histone acetylation/deacetylation is crucial in the organization of euchromatin (which enables transcription) and heterochromatin (which inhibits transcription), thereby mediating changes in gene expression. Histone deacetylases (HDACs) are members of an ancient enzyme family that reverses the acetylation of protein substrates. *HDAC*-mediated removal of acetylation from histone tail lysines generally correlates with gene silencing by decreasing the ability of transcription factors to access DNA (Chen et al. 2015). In insects, studies have confirmed that *HDACs* play a role in various developmental processes, such as growth (Ozawa et al. 2016), metamorphosis (George et al. 2019; George and Palli 2020a; George and Palli 2020b), long-term memory (Fitzsimons et al. 2013), reproduction (Zhang et al. 2018), longevity (Frankel et al. 2015), caste differentiation (Spannhoff et al. 2011; Suzuki et al. 2019), diapause (Cui et al. 2022), and immunity (Mukherjee et al. 2012). Furthermore, in the broad-horned flour beetle *Gnatocerus cornutus*, *HDAC1* and *HDAC3* differentially participate in the nutrition-dependent growth of wings and male-exaggerated mandibles, suggesting that *HDACs* may serve as epigenetic regulators linking nutritional conditions to trait-specific development (Ozawa et al. 2016). Here we expand on this work by assessing the function of diverse histone deacetylases in the structure-specific growth of both sex-shared and sex-specific traits including evolutionarily novel structures in a horned dung beetle.

Horned dung beetles (genus *Onthophagus*) have emerged as promising model systems to investigate the development and diversification of scaling relationships. Here we employ one such model, the bull-headed dung beetle *O. taurus*, to investigate the function of five different *HDAC* members in the development of four different morphological structures. We selected hind legs as examples of traits that exhibit moderate nutrition-responsiveness and therefore scale roughly isometrically with body size. We also investigated male genitalia because of their relatively muted nutrition response and corresponding hypoallometric scaling. Finally, we assessed thoracic horns and head horns because of their highly sex-specific growth and scaling relationships (Casasa et al. 2017). Thoracic horns are observed only in the pupal stage of both sexes where they function as molting devices in the shedding of the larval head capsule during the larval-pupal molt, and exhibit exaggerated growth in males (Moczek et al. 2006; Moczek 2006). In partial contrast, head horns are found only in male pupae as well as adults and exhibit extreme nutrition-responsive growth resulting in hyper-allometric scaling with body size. Head horns function as weapons in competition between adult males over reproductive access to females. While thoracic horns have recently been identified as partial wing serial homologs (Hu et al. 2019), head horns lack any obvious homology with other structures and are thus considered evolutionary novelties even by the strictest of definitions (Müller and Wagner 1991; Moczek 2005). Below we detail our results and discuss them in the light of the developmental regulation of growth and plasticity in horned beetles in particular and insects broadly.

## RESULTS

We sought to characterize the presence and function of *HDACs* during the development of the bull-headed dung beetle *O. taurus*. We identified five HDACs in the annotated genome, which, based on phylogenetic analysis, could be classified into three classes: class I (HDAC1 and HDAC3), class II (HDAC4 and HDAC6), and class IV (HDAC11) (Fig. S1). It is worth noting that a HDAC protein that was predicted as HDAC5 for *O. taurus* in NCBI (reference number: XP_022905538.1) was found to be nested within the cluster of HDAC4 proteins across insect species (Fig. S1). As a result, we have re-annotated this protein as Ot-HDAC4. We then performed RNAi experiments by injecting double-stranded RNA (dsRNA) corresponding to each of the five HDACs into newly molted final-instar *O. taurus* larvae and assessed their influence on pupal and adult morphologies. Wildtype morphology for each focal phenotype is shown in Figure 1.

**Fig. 1.**
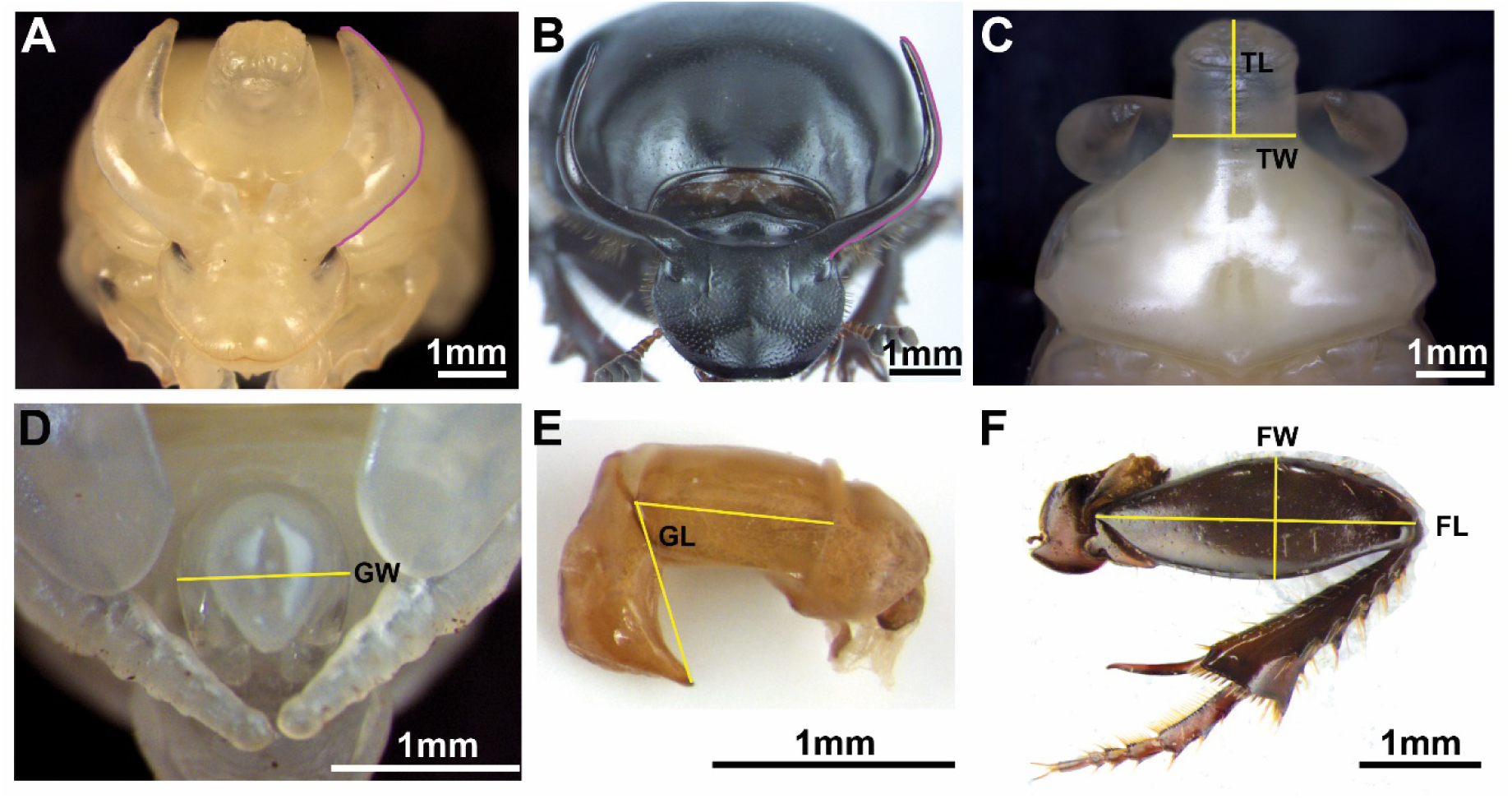
Wildtype morphology and morphometric landmarks used for morphological quantification. (**A**-**F**) Morphology of the male pupal head (A), male adult head (B), pupal pronotum from dorsal view (C), pupal reproductive organ (D), adult aedeagus (E), and hind leg (F), respectively, and the morphometric landmarks used for measurements (purple and yellow line). TW, thoracic width; TL, thoracic length; GW, genital width; GL, genital length; FL, femur length; FW, femur width.

### *HDAC*-RNAi resulted in highly variable mortality depending on the target *HDAC*

RNAi-mediated knockdown of *HDAC1* resulted in 100% larval mortality at the initial 1 μg/μl dsRNA injection dosage, as well as subsequent dosages as low as 0.25 μg/μl (Table S1). Most larvae exhibited arrested development at the pre-pupal stages and eventually died with larval-pupal intermediate phenotypes with pupal traits, such as compound eyes, observed underneath the larval integument. A mass of fat body accumulated underneath the posterior region of the developing pupal abdomen, resulting in a cavity filled with hemolymph between the larval integument and newly-formed pupal cuticle (Fig. 2A and B). When dsRNA concentration was decreased to 0.01 μg/μl, very few individuals succeeded to develop to pupal (3/44) and adult (2/44) stages (Table S1) amenable to phenotyping. In marked contrast, RNAi targeting *HDAC 3*, *4*, *6*, or *11* resulted in mortalities ranging from 16.7% to 86.7% depending on dosage and permitted more nuanced and quantitative analysis of phenotypic effects (Table S1). Because no observable phenotypes were found following *HDAC6*^RNAi^ or *HDAC11*^RNAi^ even at high dsRNA dosage, we focus on *HDAC 1*, *3*, and *4* in the remainder.

**Fig. 2.**
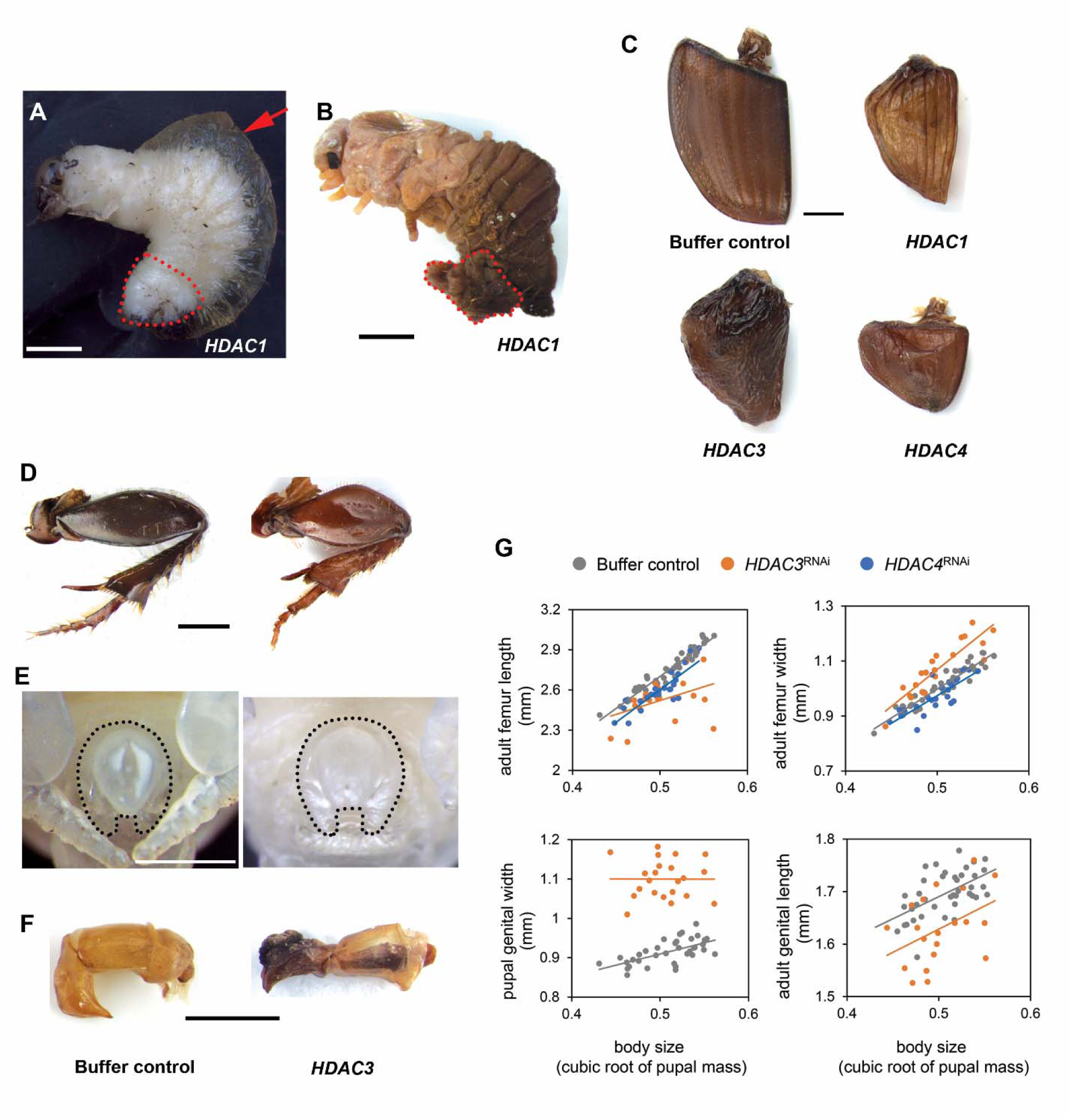
*HDAC^RNAi^* effects on molting and appendage development. (**A** and **B**) Larval-pupal intermediate induced by *HDAC1* knockdown. The fat body accumulated outside of newly-formed pupa is outlined (red dotted line) before (A) and after (B) peeling away the larval cuticle, respectively. Hemolymph in the cavity between larval and newly-formed pupal cuticle is indicated (red arrow in A). (**C**) Representative wing phenotypes are shown as follows: buffer injection, *HDAC1*^RNAi^, *HDAC3*^RNAi^, and *HDAC4*^RNAi^. (**D**-**F**) Morphology of the hind leg (D), pupal genitalia (E), and adult aedeagus (F) compared to buffer injected (left column) and *HDAC3*^RNAi^ (right column) individuals, respectively. (**G**) Knockdown of *HDAC3* (orange dots) or *HDAC4* (blue dots) on femur length and width compared to buffer-injected control (gray dots). Effects of *HDAC3*^RNAi^ on the male pupal and adult reproductive organ (bottom row in **G**). The RNAi phenotypes and their corresponding negative controls are shown at the same magnification. Scale bars: 1 mm.

### *HDACs* function during appendage development

To determine affected traits, we measured the scaling relationship of each trait to the cubic root of pupal mass as a proxy of body size since the more commonly used measure of thorax width was affected by *HDAC3* knockdown (treatment: *P* < 0.001, Table S2). *HDAC1^RNAi^*resulted in curtailment of both forewings (i.e. elytra) and hindwings at the pupal stage, which was retained into the adult (Fig. 2C). Similarly, both *HDAC3* or *HDAC4* knockdown diminished wing size (Fig. 2C). Thus, *HDAC 1, 3,* and *4* appear to regulate wing development in similar ways. RNAi targeting *HDAC3* and *HDAC4* also affected leg development, but specific effects diverged. To assess leg phenotypes quantitatively we selected the femur, which is especially amenable to width and length measurement, for morphometric analyses. *HDAC4*^RNAi^ led to a reduction in femur length (treatment: *P* < 0.001, Table S2) while the slope of the body size/femur length allometry was decreased in *HDAC3*^RNAi^ animals (treatment: *P* = 0.163; treatment × body size: *P* = 0.004, Fig. 2D and G, and Table S2). In contrast, whereas *HDAC3*^RNAi^ increased femur width compared to control individuals (treatment: *P* < 0.001, Table S2), the same measure was reduced in *HDAC4* knockdown animals (treatment: *P* = 0.001, Fig. 2D and G, and Table S2). Lastly, we found that knockdown of *HDAC3* also affected the development of the male reproductive organ, the aedeagus, itself composed of the more proximal phallobase and the more distal parameres. Specifically, genital width increased and attained a more rounded shape at the pupal stage (treatment: *P* < 0.001, Fig. E and G, and Table S2), while adult genitalia exhibited a deformation and overall shortening of both parameres and phallobase, which combined yielded a shortening of overall aedeagus length across all body sizes (treatment: *P* < 0.001, Fig. F and G, and Table S2).

### *HDACs* function during thoracic and head horn formation

Head and thoracic horns are textbook examples of evolutionary novelties, and we sought to determine whether *HDAC* function may have been co-opted during the evolution of one or both horn types. *HDAC1*^RNAi^ resulted in a reduction in thoracic horn length and a split tip at the pupal stage (Fig. 3A and C), whereas the corresponding area in the adult exhibited a broad indentation (compared to the smoothly convex outline observed in wildtype or buffer control injected individuals) and small bilateral projections at the respective edge of the indentation (Fig. 3F and G). Effects on head horns could not be quantified with certainty due to the high degree of natural variability of the trait and the very low number of surviving males, which in addition were too small to develop fully formed head horns. However, *HDAC3* knockdown caused measurable shape and scaling changes in thoracic horns. Specifically, *HDAC3*^RNAi^ increased pupal thoracic horn width (treatment: *P* < 0.001, Fig. 3D, E, L, and Table S3), but decreased pupal thoracic horn length (treatment: *P* < 0.001, Fig. 3L and Table S3). In addition, we found a bilateral indentation to the distal region of the thoracic horn, causing the thoracic horns of large *HDAC3*^RNAi^ pupae to attain a more conical shape (Fig. 3B). In contrast to thoracic horn phenotypes, *HDAC3*^RNAi^ yielded drastically enlarged head horns, in particular concerning head horn *width* across the entire range of male body sizes (Fig. 3E, H-K, and Fig. S2). However, due to the highly varied nature of these phenotypes, we were unable to arrive at reliable landmarks for quantitative measure, hence this observation could only be made qualitatively. To determine whether head horn *length* was also affected, we further measured the scaling relationship of head horn length to body size. *HDAC3*^RNAi^ steepened the slope of the head horn length allometry at both pupal (treatment: *P* = 0.025, Fig. 3L and Table S3) and adult stages (treatment: *P* = 0.012, Fig. 3L and Table S3). Notably, *HDAC3*^RNAi^ also reduced the maximum asymptotic horn length in adults (treatment: *P* < 0.001, Fig. 3L and Table S3), which was not detected at the pupal stage (treatment: *P* = 0.107, Fig. 3L and Table S3). The quantification of head horn length reduction from pupa to adult further confirmed this observation (treatment: *P* < 0.05, Table S2), and this effect was enhanced with increasing body size (treatment × body size effect: *P* = 0.010, Fig. 3L and Table S2). These results suggest that *HDAC3*^RNAi^ affects head horn *width* during the horn growth phase (which takes place during the larval–to–pupal transition, thereby resulting in visible *pupal* phenotypes), but affects horn *length* during the pupal remodeling phase of ontogeny (thus becoming apparent in adults only). *HDAC4*^RNAi^ in turn not only decreased maximum asymptotic horn length in adults (treatment: *P* = 0.027, Table S3), but also altered the threshold body size of sigmoidal allometry separating small hornless from large and fully horned males, such that large head horns now also presented in males of relatively small body sizes (treatment: *P* < 0.001, Fig. 3L and Table S3). In contrast, we did not find abnormal phenotypes with respect to thoracic horns in *HDAC4*^RNAi^ individuals.

**Fig. 3.**
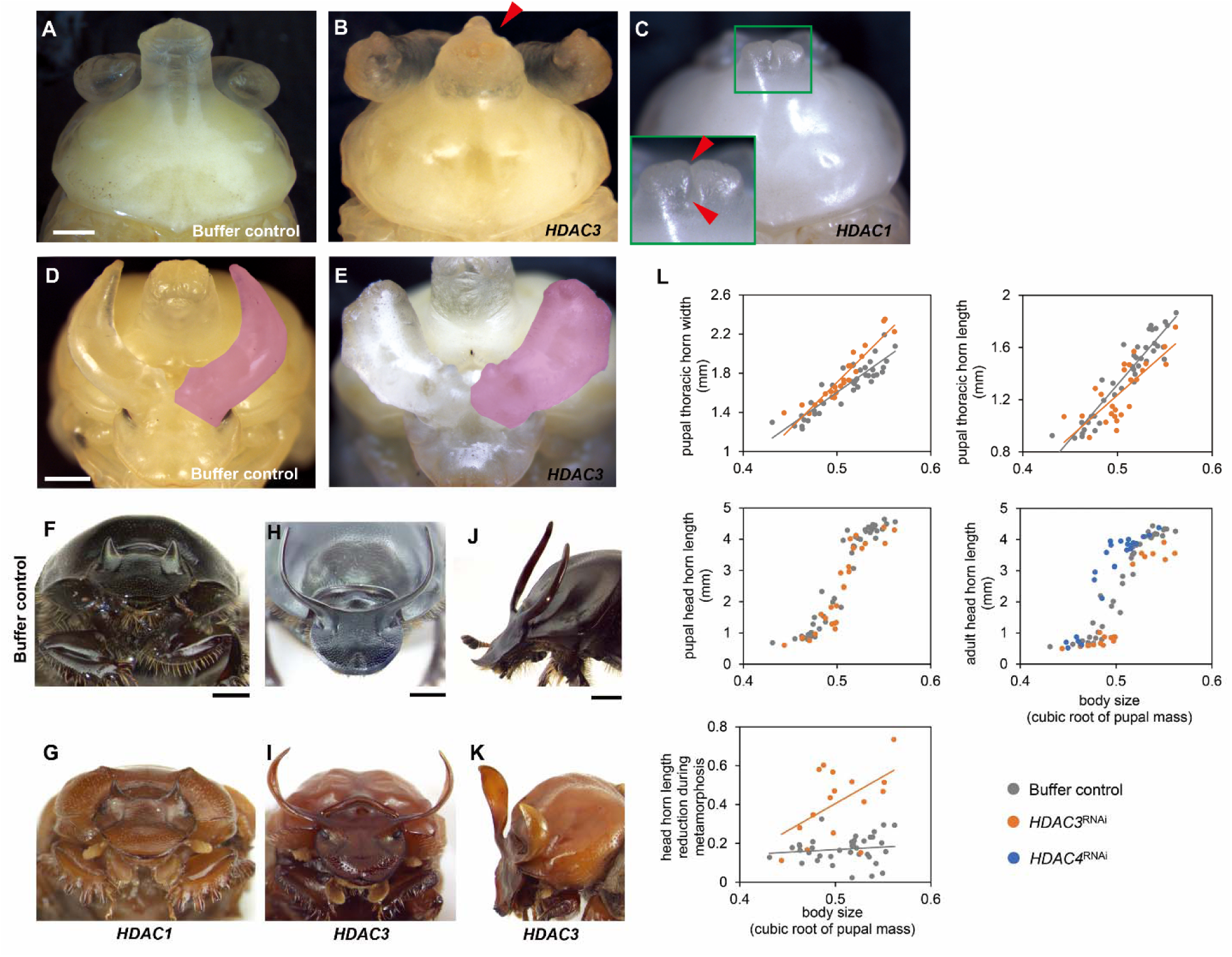
***HDAC^RNAi^* effects on thoracic and head horn formation.** (**A**-**C**) The pronotum in buffer injected (A), *HDAC3*^RNAi^ (B), and *HDAC1*^RNAi^ (C) individuals. Inset in C shows the prothoracic horn. The furrow between paired horn vestiges is indicated by red arrowheads. (**D** and **E**) Representative head horn of negative control (D) and *HDAC3*^RNAi^ (E) individuals, respectively. The right head horn is colored magenta. (**F**-**K**) The morphology of the adult pronotum in front view (F and G), front view of head horns (H and I), as well as head horn viewed from lateral (J and K) in negative control (third row) individuals and following *HDAC* ^RNAi^ (bottom row), respectively. (**L**) Changes in thoracic and head horns resulting from *HDAC3*^RNAi^ and *HDAC4*^RNAi^. Reduction in head horn length during the transition from pupa to adult is increased by *HDAC3*^RNAi^ compared to buffer-injections (measured as the difference between pupal head horn length and adult head horn length in the same individual). RNAi phenotypes and their corresponding negative controls are shown at the same magnification. Scale bars: 1 mm.

## DISCUSSION

### The significance of *HDACs* in horned beetle development and evolution

Earlier work on the broad-horned flour beetle *Gnatoceros cornutus* was the first to document the role of *HDACs* in the regulation of nutrition-responsive plasticity in insects (Ozawa et al. 2016). Large males in this species develop conspicuous mandibular projections (called mandibular horns) which were reduced following *HDAC1*^RNAi^, whereas *HDAC3*^RNAi^ led to hypertrophy. Opposite effects were observed with respect to wing size, yet none in genitalia (Ozawa et al. 2016). These results were the first to suggest that *HDACs* operate in a trait-specific manner, and in particular contribute to the plastic and sex-specific expression of exaggerated male mandibles. Our results presented here further support a role of *HDACs* in the regulation of trait-specific plasticity, as well as add important new aspects to our understanding of HDAC function in insect development.

First, similar to the horns of rhinoceros beetle *Trypoxylus dichotomus*, *Onthophagus* horns frequently exhibit pronounced nutrition-dependent plasticity, in contrast to the more isometric growth typical of wings and legs, or the hypo-allometric growth of genitalia (Emlen et al. 2012; Casasa et al. 2017). Several mechanisms have been proposed as possibly underlying module-specific conditional growth (e.g. insulin/insulin-like growth factor (Emlen et al. 2012; Okada et al. 2019), FOXO (Tang et al. 2011; Snell-Rood and Moczek 2012; Casasa and Moczek 2018), HDAC (Ozawa et al. 2016)). Ozawa *et al*. (2016), in particular, proposed a mechanistic explanation of epigenetic flexibility in which developmentally plastic organs (e.g. mandibular horns in *G. cornutus*) are more susceptible to epigenetic (i.e. *HDAC*) perturbation, whereas developmentally robust organs (e.g. genitalia) are non-responsive to *HDAC* perturbation. However, results presented here are at odds with this model. Specifically, even though head and thoracic horn development in *O. taurus* exhibit exaggerated nutritional plasticity, the effects of *HDAC3*^RNAi^ were considerably more pronounced in genitalia and wings.

Genitalia in particular exhibited a considerable reduction of size relative to body size across all body sizes following *HDAC3*^RNAi^, thereby highlighting a previously unexpected role of *HDAC* in regulating the development of traits generally assumed to be robust to nutritional variation. Intriguingly, a similar outcome was observed following *insulin receptor* (*InR1*/*2*) transcript depletion in *O. taurus* (Casasa and Moczek 2018). Hence, the epigenetic flexibility hypothesis proposed in *Gnatoceros* beetles is unlikely to explain the findings seen here in *Onthophagus*, consistent with divergences in *HDAC* function across the Coleoptera, again similar to what has recently been reported for the insulin signaling pathway (Rohner et al. 2023). This in turn raises the possibility that, in addition to its primary function in regulating epigenetic status, *HDAC3* may also function in aspects of trait morphogenesis not related to nutritional status and developmental plasticity.

Second, we found that *HDAC1*^RNAi^ induced developmental arrest at the prepupal stages in line with previous studies in *Tribolium castaneum* (George et al. 2019), which suggests that *HDAC1* expression is required for suppressing the juvenile hormone (JH) response gene *Krüppel homolog 1* (*Kr-h1*). In *Tribolium*, *HDAC1* knockdown prevents larval-to-pupal transition via derepressing *Kr-h1* expression, thereby influencing JH actions and thus halting metamorphosis. Similarly, high dose of *HDAC1* knockdown caused developmental failures during the pupal stage in *Gnatoceros*, indicating a possibly conserved role in basic developmental process mediating metamorphosis. However, despite this putative conservation of *HDAC1* function across the Coleoptera assessed to date, we also found that the precise nature and direction of *HDAC*^RNAi^ effects on appendage formation diverged between *Gnatoceros* and *Onthophagus* beetles even beyond those already noted above: for example, in *Onthophagus* downregulation of *HDAC1*, *3*, and *4* all appear to affect wing size similarly, whereas in *Gnatoceros HDAC1*^RNAi^ and *HDAC3*^RNAi^ yield opposite effects. Likewise, in *Onthophagus*, *HDAC3*^RNAi^ increased femur width, but *HDAC4*^RNAi^ decreased it, whereas leg morphology was generally unaffected in *Gnatoceros* beetle. Lastly, our results document the recruitment of *HDAC* function into the formation of an evolutionarily novel structure - head and thoracic horns - suggesting that *HDAC* function is not just evolutionarily labile among conserved insect traits but also contributed to the comparatively recent evolution of *Onthophagus* weaponry, including the regulation of size, shape, and key components of scaling.

### Development and evolution of pupal remodeling

The horns of adult beetles are the product of developmental processes operating at at least two distinct stages of development, a rapid growth phase approximately 48h immediately prior to the larval-to-pupal molt and a remodeling phase during the pupal stage (Moczek 2006). While generally given less attention, pupal remodeling can be quite extensive and fully formed pupal horns may be subject to considerable reduction and even complete resorption in many species. Thus, the morphological diversity of adult horns is not only influenced by the differential regulation of growth during the prepupal stage, but also by the developmental processes underlying the differential resorption of horn tissue during the pupal stage (Moczek 2006; Moczek 2007; Morita et al. 2019). Previous work identified that differential programmed cell death facilitates species, sex, and body-region specific resorption of horn primordia (Kijimoto et al. 2010). However, the mechanisms regulating horn resorption during the pupal stage remain largely unknown. Our results implicate *HDAC3* as a regulator of both prepupal growth and pupal remodeling of horn primordia. Specifically, we show that *HDAC3*^RNAi^ increased head horn width, that this phenotype was already prominently visible at the pupal stage, and must therefore have resulted from modifications to the prepupal growth phase of horn formation (Fig. 3). In addition, however, we also find that *HDAC3*^RNAi^ altered horn *length* in a manner not evident at the pupal stage but clearly discernible in the resulting adults, and thus a consequence of *HDAC3*^RNAi^ effects on the pupal remodeling phase of horn formation. As such, *HDAC3* is one of relatively few genes identified to date to be involved in horn remodeling during the pupal stage of horned beetles (Morita et al. 2019; Matsuda et al. 2023).

### Chromatin modifications and developmental plasticity

This work is the first to implicate chromatin modifications in the regulation of development and plasticity in horned dung beetles. While earlier work documented the existence of the complete methylation machinery in the *O. taurus* genome alongside sex- and nutrition-dependent differences in methylation signatures, the functional significance of chromatin modifications, if any, had remained unknown (Choi et al. 2010; Snell-Rood and Moczek 2013). We now show that downregulation of *HDAC3* and *HDAC4* affect critical aspects of horn formation including size, shape, and the location of the inflection point separating alternate male morphs. Future work will need to explore if and how *HDAC* functions may be contributing to the genome-wide remodeling patterns, and more generally the roles of *cis*-regulatory elements in the development and evolution of plasticity in insects.

## METHODS

### Insects

Adult *O. taurus* were collected, courtesy of John Allen, from Paterson Farm near Ravenswood, Western Australia. A laboratory population was maintained at 25°C in a sand/soil mixture and fed cow manure twice a week. Larvae used for injection were collected and prepared as described previously (Shafiei et al. 2001).

### Identification of *O. taurus* orthologs for HDACs

The *Onthophagus* orthologs of *HDAC* genes were identified via reciprocal BLAST to *T. castaneum*, *G. cornutus*, *Drosophila melanogaster*, and *Homo sapiens* in NCBI databases. Amino acid sequences of *HDAC* genes from the above species were aligned with ClustalW algorithm implemented in MEGA X (Kumar et al. 2018). Neighbor-Joining trees were constructed using MEGA X.

### Double-stranded RNA (dsRNA) synthesis and injection

Total RNA was extracted with RNeasy Mini Plus Kit (QIAGEN) and reverse transcribed with iScript cDNA Synthesis kit (Bio-Rad). Partial fragments of each genes were amplified with PCR by using gene-specific primers and cloned into pCR4-TOPO TA vector (Invitrogen, Thermo Fisher Scientific). DNA templates for *in vitro* transcription were produced with PCR by using TOPO RNAi primer (Supplementary table S4) (Philip and Tomoyasu 2011). PCR products were purified and concentrated using the QIAquick PCR Purification kit (QIAGEN) and subjected to *in vitro* transcription (MEGAscript T7 Transcription Kit, Thermo Fisher Scientific) and dsRNA purification (MEGAclear Transcription Clean-Up Kit, Thermo Fisher Scientific), according to the manufacturer’s protocol. DsRNA was quantified and stored at -80°C until use. Injection was carried out at the early stage of the last larval instar (i.e. the third larval instar, L3). Control animals were injected with injection buffer (1.4 mM NaCl, 0.07mM Na_2_HPO_4_, 0.03mM KH_2_PO_4_, and 4mM KCl) kept at the same condition as dsRNA injected animals (See Supplementary table S1 for detailed injection information).

### Effect of *HDAC3*- and *HDAC4*-RNAi on the scaling relationships between several morphological traits and body size

Since the usual measure of *Onthophagus* body size - thorax width (e.g. Casasa and Moczek 2018; Macagno et al. 2021) - was clearly affected by our RNAi treatments (Fig. 3B and Table S2), we measured the cube root of pupal mass as a proxy for individual body size (Marchini et al. 2014; Fleming et al. 2021). We used t-tests to compare body sizes between each RNAi-treated and control group.

Consistent with previous studies we analyzed nonlinear horn allometries using untransformed data (e.g. Casasa and Moczek 2018; Macagno et al. 2021). We used the package drc (Ritz et al. 2015) in R 3.5.2 (R Core Team 2018) to fit the body size/head horn length distribution a four-parameter log-logistic (Hill) function in the form

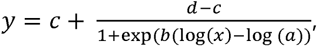

with *x* = body size, *y* = head horn length, *a* = body size at the point of inflection of the sigmoid curve, *b* = slope of the curve, *c =* minimum and *d* = maximum asymptotic horn lengths (Macagno et al. 2021). We then inferred whether a complex model including a sigmoidal regression per treatment (i.e. control and RNAi treatments) fitted our data better than a simpler model with one sigmoidal regression including the whole sample by means of the Akaike Information Criterion (AIC) (Akaike 1974). The AIC measures relative model fit - the lower its value, the better the model fits to the experimental data (Burnham et al. 2011). Upon finding the complex model more fitting, we used Welch’s t-tests (with Holm-Bonferroni sequential correction where applicable) to compare parameter means (*a*, *b*, *c*, *d*) between control-injected and RNAi treatment groups (Moczek et al. 2002; Casasa and Moczek 2018). We compared control individuals to *HDAC3*^RNAi^ and *HDAC4*^RNAi^ individuals in the case of the adult horn allometry. As for pupal horn allometry, we compared *HDAC3* ^RNAi^ to control individuals.

To inspect the effect of RNAi manipulations on all other morphological traits, we modeled trait size as a function of body size, treatment (*HDAC3*^RNAi^ or *HDAC4*^RNAi^ vs control-injected) and their interaction using general linear models in SPSS Statistics 25 (IBM Corp., 2017). Models were simplified by removing non-significant interactions. Data were log-transformed prior to analyses (Huxley 1924). The reduction in horn size during metamorphosis (defined as pupal horn length - adult horn length) was analyzed similarly with a general linear model, using untransformed data as in other analyses of horn morphology. Statistics for morphometric analyses are provided in Supplementary Table S2 and S3.

### Image Processing

All images were captured with a digital camera (Scion) mounted to a dissecting microscope (Leica MZ16, Germany). Brightness and contrast of images were adjusted across the entire image with Adobe Photoshop CC 2017 (Adobe, USA).

## DECLARATIONS

### Ethical Approval

Not applicable.

### Availability of data and materials

All data are available in the manuscript and supplementary materials.

### Authors’ contributions

Y.H. conceived the initial idea, Y.H. and A.P.M. designed the research, Y.H. and J.R.C. performed the experiments, Y.H. and A.L.M.M. analyzed the data, Y.H., A.P.M., and A.L.M.M. wrote the manuscript with contribution from J.R.C.

## Acknowledgements

We would like to thank Kayla Copper and Levi Burdine for help with beetle care, and Patrick Rohner for discussion of statistical analyses.

