## Supplementary material S1 for "Histone deacetylases regulate organ-specific growth in a horned beetle" (from another version of this preprint: version 1, 2023-12-01)

**This file includes:**

Figs. S1 to S2
Tables S1 to S4

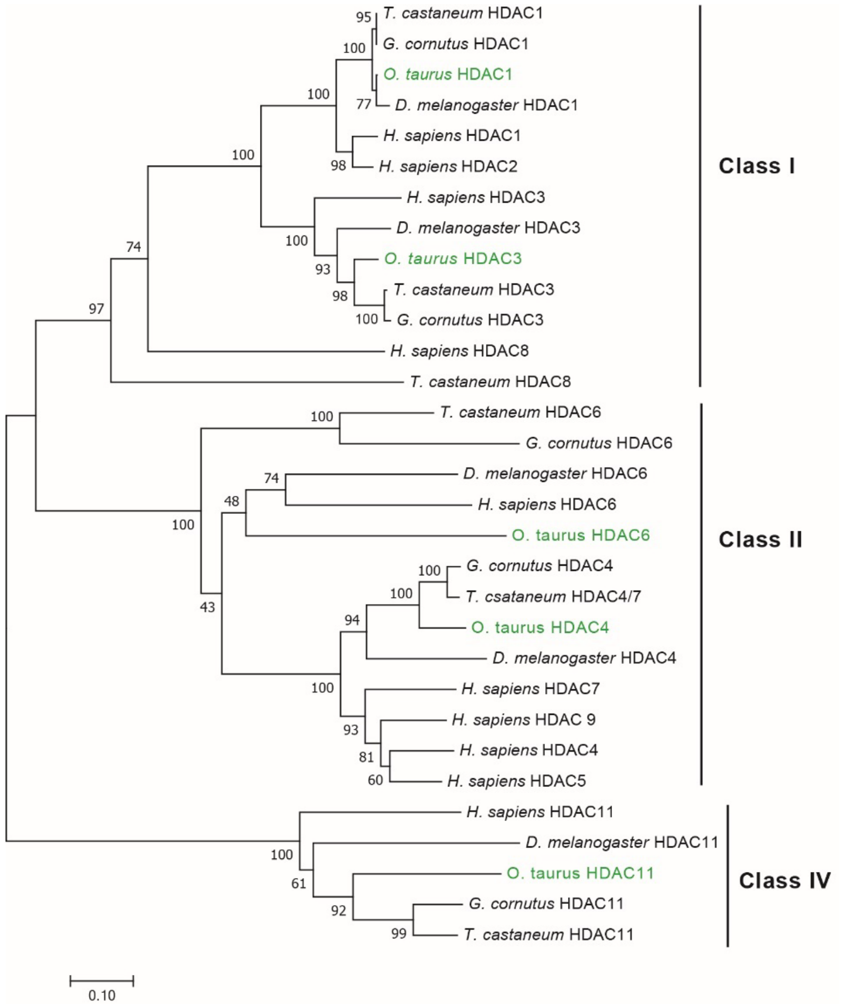


**Fig. S1. Phylogenetic analysis of HDACs**

Protein sequences of HDACs were aligned and then subjected to construction of Neighbor-joining phylogenetic tree. The NCBI accession numbers of protein sequences are listed as follows: *T. castaneum* HDAC1 (XP_966633.1), *G. cornutus* HDAC1 (BAW19561.1), *O. taurus* HDAC1 (XP_022902140.1), *D. melanogaster* HDAC1 (NP_647918.2), *H. sapiens* HDAC1 (BAG70111.1), *H. sapiens* HDAC2 (NP_001518.3), *T. castaneum* HDAC3 (XP_969419.1), *G. cornutus* HDAC3 (BAW19562.1), *O. taurus* HDAC3 (XP_022899704.1), *D. melanogaster* HDAC3 (NP_651978.2), *H. sapiens* HDAC3 (AAC52038.1), *H. sapiens* HDAC8 (AAF73428.1), *T. castaneum* HDAC8 (XP_008192051.1), *T. castaneum* HDAC6 (EFA11522.2), *G. cornutus* HDAC6 (BAW19564.1), *D. melanogaster* HDAC6 (ACN94041.1), *H. sapiens* HDAC6 (AAD29048.1), *O. taurus* HDAC6 (XP_022914414.1), *G. cornutus* HDAC4 (BAW19563.1), *T. castaneum* HDAC4/7 (XP_967425.2), *O. taurus* HDAC4 (XP_022905538.1), *D. melanogaster* HDAC4 (NP_001259507.1), *H. sapiens* HDAC7 (CAB55935.1), *H. sapiens* HDAC9 (AAO27363.1), *H. sapiens* HDAC4 (NP_001365345.1), *H. sapiens* HDAC5 (KAI2583350.1), *H. sapiens* HDAC11 (NP_079103.2), *D. melanogaster* HDAC11 (NP_001247296.1), *O. taurus* HDAC11 (XP_022906835.1), *G. cornutus* HDAC11 (BAW19565.1), *T. castaneum* HDAC11 (XP_970656.1).

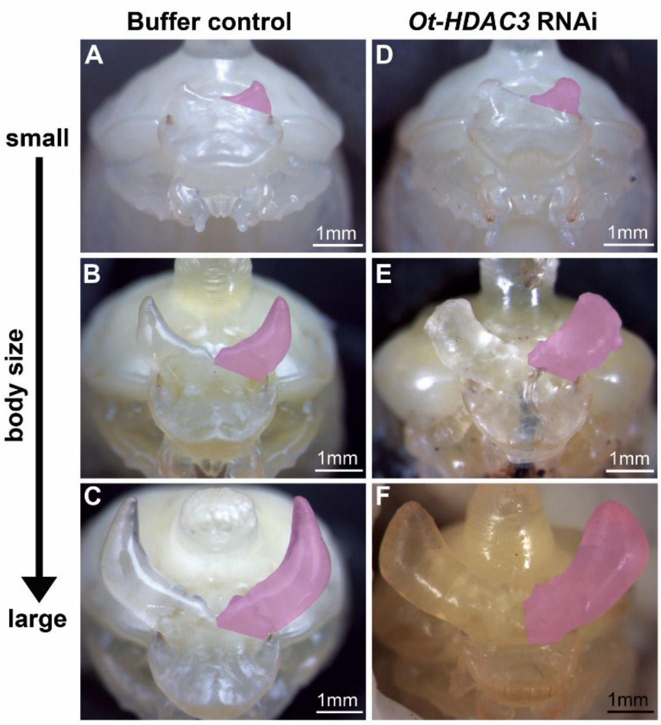


**Fig. S2.** ***HDAC3*$^{RNAi}$ head horn phenotypes across the range of male body sizes**

(**A**-**F**) Dorsal view of the pupal head in buffer injected control individuals (left column) compared to *HDAC3*$^{RNAi}$ individuals (right column) in (from top to bottom) small, medium-sized and large individuals, respectively. The right head horn is colored magenta.

### Table S1. Injection information for this study

| dsRNA | concentration (µg/µl) | number of injected animals | survived to Pupa | survived to adult | mortality (died before eclosion) | phenotypic penetrance |
|---|---|---|---|---|---|---|
| ***HDAC1*** | 1.0 | 25 | 0 | 0 | 100% | 100% |
| | 0.5 | 22 | 0 | 0 | 100% | 100% |
| | 0.25 | 53 | 0 | 0 | 100% | 100% |
| | 0.1 | 71 | 1 | 0 | 100% | 100% |
| | 0.05 | 54 | 4 | 3 | 94.4% | 50% |
| | 0.01 | 44 | 3 | 2 | 95.5% | 66.6% |
| ***HDAC3*** | 0.5 | 176 | 43 | 29 | 83.4% | 95.3% |
| | 1.0 | 60 | 12 | 8 | 86.7% | 100% |
| ***HDAC4*** | 0.5 | 62 | 39 | 35 | 43.5% | 89.7% |
| | 1.0 | 18 | 7 | 7 | 61.1% | 100% |
| ***HDAC6*** | 1.0 | 18 | 16 | 15 | 16.7% | 0% |
| | 2.0 | 23 | 20 | 19 | 17.4% | 0% |
| ***HDAC11*** | 2.0 | 18 | 11 | 11 | 38.9% | 0% |
| | 1.0 | 30 | 25 | 24 | 20% | 0% |
| **buffer** | N/A | 62 | 57 | 56 | 9.7% | 0% |

### Table S2. The primer information used in this study

| name | primer F | primer R | GenBank Accession No. | length (bp) | purpose |
|---|---|---|---|---|---|
| *HDAC1* | CGTCAAACGTTACGG GCTTC | GCGATTGTCCCTCC TTCCAC | XM_023046372.1 | 445 | gene clone |
| *HDAC3* | TGTTTTGCAATGTGGG GCTG | GAATCGCGTCTGGT GGGATA | XM_023043936.1 | 401 | gene clone |
| *HDAC4* | ACCTTGCGGAGGAAT TGGAG | CGAATCCAGCGGAA ACCAAC | XM_023049770.1 | 554 | gene clone |
| *HDAC6* | GTTGGATGATGCACC AGGGA | AATCATAGCCACAC CCGCTC | XM_023058646.1 | 643 | gene clone |
| *HDAC11* | GGCCCATTATCTATAG GGACG | AGTGATGTCAGCAT AAGGGCA | XM_023051067.1 | 431 | gene clone |
| TOPO RNAi Primer | TAATACGACTCACTAT AGGGCGAATTCGCCC TT | TAATACGACTCACTA TAGGGCGAATTCGC CCTT | N/A | N/A | DNA template for *in vitro* transcription |

### Table S3. Effects of *HDAC3* and *HDAC4* knockdowns on trait sizes

| | Type III Sum of Squares | df | Mean Square | *F* | *P* |
|---|---|---|---|---|---|
| Effect of *HDAC3*-RNAi on thoracic width | | | | | |
| Corrected Model | 0.202[a] | 2 | 0.101 | 111.240 | 0.000 |
| Intercept | 13.652 | 1 | 13.652 | 15059.789 | 0.000 |
| Treatment | 0.017 | 1 | 0.017 | 18.594 | 0.000 |
| ln_curtPupaMass | 0.191 | 1 | 0.191 | 210.328 | 0.000 |
| Error | 0.056 | 62 | 0.001 | | |
| Total | 485.845 | 65 | | | |
| Corrected Total | 0.258 | 64 | | | |
| a. $R^2$ = .782 (Adjusted $R^2$ = .775) | | | | | |
| Effect of *HDAC3*-RNAi on thoracic length | | | | | |
| Corrected Model | 0.599[a] | 2 | 0.300 | 502.653 | 0.000 |
| Intercept | 10.053 | 1 | 10.053 | 16866.410 | 0.000 |
| Treatment | 0.089 | 1 | 0.089 | 148.607 | 0.000 |
| ln_curtPupaMass | 0.485 | 1 | 0.485 | 813.509 | 0.000 |
| Error | 0.037 | 62 | 0.001 | | |
| Total | 429.138 | 65 | | | |
| Corrected Total | 0.636 | 64 | | | |
| a. $R^2$ = 0.942 (Adjusted $R^2$ = 0.940) | | | | | |
| Effect of *HDAC3*-RNAi on adult femur length | | | | | |
| Corrected Model | 0.234[a] | 3 | 0.078 | 65.738 | 0.000 |
| Intercept | 6.603 | 1 | 6.603 | 5555.831 | 0.000 |
| Treatment | 0.002 | 1 | 0.002 | 1.992 | 0.163 |
| ln_curtPupaMass | 0.091 | 1 | 0.091 | 76.322 | 0.000 |
| Treatment × ln_curtPupaMass | 0.011 | 1 | 0.011 | 9.003 | 0.004 |
| Error | 0.071 | 60 | 0.001 | | |
| Total | 251.072 | 64 | | | |
| Corrected Total | 0.306 | 63 | | | |
| a. $R^2$ = 0.767 (Adjusted $R^2$ = 0.755) | | | | | |
| Effect of *HDAC4*-RNAi on adult femur length | | | | | |
| Corrected Model | 0.221[a] | 2 | 0.111 | 268.207 | 0.000 |
| Intercept | 6.694 | 1 | 6.694 | 16248.194 | 0.000 |
| Treatment | 0.022 | 1 | 0.022 | 53.876 | 0.000 |
| ln_curtPupaMass | 0.181 | 1 | 0.181 | 438.427 | 0.000 |
| Error | 0.027 | 65 | 0.000 | | |
| Total | 268.622 | 68 | | | |
| Corrected Total | 0.248 | 67 | | | |
| a. $R^2$ = 0.892 (Adjusted $R^2$ = 0.889) | | | | | |
| Effect of *HDAC3*-RNAi on adult femur width | | | | | |
| Corrected Model | 0.360[a] | 2 | 0.180 | 205.228 | 0.000 |
| Intercept | 1.143 | 1 | 1.143 | 1301.267 | 0.000 |

| | | | | | |
|---|---|---|---|---|---|
| Treatment | 0.064 | 1 | 0.064 | 72.840 | 0.000 |
| ln_curtPupaMass | 0.318 | 1 | 0.318 | 361.927 | 0.000 |
| Error | 0.054 | 61 | 0.001 | | |
| Total | 68.016 | 64 | | | |
| Corrected Total | 0.414 | 63 | | | |
| a. $R^2$ = 0.871 (Adjusted $R^2$ = 0.866) | | | | | |
| Effect of *HDAC4*-RNAi on adult femur width | | | | | |
| Corrected Model | 0.279[a] | 2 | 0.140 | 177.141 | 0.000 |
| Intercept | 1.017 | 1 | 1.017 | 1289.299 | 0.000 |
| Treatment | 0.009 | 1 | 0.009 | 11.774 | 0.001 |
| ln_curtPupaMass | 0.255 | 1 | 0.255 | 322.847 | 0.000 |
| Error | 0.051 | 65 | 0.001 | | |
| Total | 67.823 | 68 | | | |
| Corrected Total | 0.331 | 67 | | | |
| a. $R^2$ = 0.845 (Adjusted $R^2$ = 0.840) | | | | | |
| Effect of *HDAC3*-RNAi on pupal genital width | | | | | |
| Corrected Model | 0.471[a] | 2 | 0.236 | 183.235 | 0.000 |
| Intercept | 1.876 | 1 | 1.876 | 1458.133 | 0.000 |
| Treatment | 0.468 | 1 | 0.468 | 364.017 | 0.000 |
| ln_curtPupaMass | 0.011 | 1 | 0.011 | 8.288 | 0.006 |
| Error | 0.071 | 55 | 0.001 | | |
| Total | 56.279 | 58 | | | |
| Corrected Total | 0.542 | 57 | | | |
| a. $R^2$ = 0.870 (Adjusted $R^2$ = 0.865) | | | | | |
| Effect of *HDAC3*-RNAi on adult genital length | | | | | |
| Corrected Model | 0.036[a] | 2 | 0.018 | 23.366 | 0.000 |
| Intercept | 4.164 | 1 | 4.164 | 5446.965 | 0.000 |
| Treatment | 0.019 | 1 | 0.019 | 24.384 | 0.000 |
| ln_curtPupaMass | 0.014 | 1 | 0.014 | 18.354 | 0.000 |
| Error | 0.044 | 57 | 0.001 | | |
| Total | 137.709 | 60 | | | |
| Corrected Total | 0.079 | 59 | | | |
| a. $R^2$ = 0.451 (Adjusted $R^2$ = 0.431) | | | | | |
| Effect of *HDAC3*-RNAi on pupal thoracic length | | | | | |
| Corrected Model | 2.275[a] | 2 | 1.138 | 120.467 | 0.000 |
| Intercept | 0.210 | 1 | 0.210 | 22.221 | 0.000 |
| Treatment | 0.060 | 1 | 0.060 | 6.307 | 0.015 |
| ln_curtPupaMass | 2.167 | 1 | 2.167 | 229.471 | 0.000 |
| Error | 0.586 | 62 | 0.009 | | |
| Total | 108.091 | 65 | | | |
| Corrected Total | 2.861 | 64 | | | |
| a. $R^2$ = 0.795 (Adjusted $R^2$ = 0.789) | | | | | |
| Effect of *HDAC3*-RNAi on pupal thoracic width | | | | | |
| Corrected Model | 1.271[a] | 2 | 0.635 | 171.029 | 0.000 |

| | | | | | |
|---|---|---|---|---|---|
| Intercept | 1.460 | 1 | 1.460 | 393.029 | 0.000 |
| Treatment | 0.070 | 1 | 0.070 | 18.959 | 0.000 |
| ln_curtPupaMass | 1.230 | 1 | 1.230 | 331.062 | 0.000 |
| Error | 0.230 | 62 | 0.004 | | |
| Total | 151.295 | 65 | | | |
| Corrected Total | 1.501 | 64 | | | |
| a. $R^2$ = 0.847 (Adjusted $R^2$ = 0.842) | | | | | |
| Effect of *HDAC3*-RNAi on head horn reduction during metamorphosis | | | | | |
| Corrected Model | 0.796[a] | 3 | 0.265 | 25.009 | 0.000 |
| Intercept | 0.043 | 1 | 0.043 | 4.095 | 0.048 |
| Treatment | 0.050 | 1 | 0.050 | 4.666 | 0.036 |
| curtPupaMass | 0.113 | 1 | 0.113 | 10.665 | 0.002 |
| Treatment × curtPupaMass | 0.077 | 1 | 0.077 | 7.236 | 0.010 |
| Error | 0.520 | 49 | 0.011 | | |
| Total | 4.461 | 53 | | | |
| Corrected Total | 1.316 | 52 | | | |
| a. $R^2$ = 0.605 (Adjusted $R^2$ = 0.581) | | | | | |

Trait size was modeled as a function of body size, treatment (*HDAC3*-RNAi or *HDAC4*-RNAi vs control-injected) and their interaction using general linear models. Models were simplified by removing non-significant interactions ($P > 0.05$).

**Table S4. Effects of *HDAC3* and *HDAC4* knockdowns on thoracic and head horns**

| | Buffer injection | | | | *HDAC3* | | | | Buffer VS *HDAC3* |
|---|---|---|---|---|---|---|---|---|---|
| | estimate | SE | SD | N | estimate | SE | SD | N | *P* |
| slope | -43.130 | 7.824 | 53.635 | 47 | -83.928 | 19.671 | 98.353 | 25 | 0.0254 |
| lower | 0.748 | 0.177 | 1.210 | 47 | 0.982 | 0.165 | 0.823 | 25 | 0.39 |
| upper | 4.506 | 0.143 | 0.979 | 47 | 4.128 | 0.169 | 0.845 | 25 | 0.107 |
| inflection | 0.501 | 0.002 | 0.016 | 47 | 0.506 | 0.002 | 0.008 | 25 | 0.1475 |

| | Buffer | | | | *HDAC4* | | | | HDAC3 | | | | Buffer VS *HDAC4* | Buffer VS *HDAC3* |
|---|---|---|---|---|---|---|---|---|---|---|---|---|---|---|
| | estimate | SE | SD | N | estimate | SE | SD | N | estimate | SE | SD | N | *P* | *P* |
| slope | -44.315 | 6.865 | 44.488 | 42 | -49.251 | 11.971 | 59.855 | 25 | -118.660 | 39.519 | 176.734 | 20 | 0.701 | 0.0123 |
| lower | 0.656 | 0.139 | 0.901 | 42 | 0.254 | 0.319 | 1.597 | 25 | 0.665 | 0.102 | 0.454 | 20 | 0.1915 | 0.9666 |
| upper | 4.403 | 0.128 | 0.827 | 42 | 3.971 | 0.123 | 0.616 | 25 | 3.578 | 0.127 | 0.568 | 20 | 0.027 | 0.0002 |
| inflection | 0.505 | 0.002 | 0.012 | 42 | 0.475 | 0.003 | 0.013 | 25 | 0.509 | 0.003 | 0.013 | 20 | 0.0001 | 0.237 |

Model fitted: Log-logistic (ED50 as parameter) (4 parms). Slope = slope of each sigmoid (b), lower = minimum asymptotic horn length (c), upper = maximum asymptotic horn length (d), inflection = body size at the inflection point (a).
